# Removing auto-activators from yeast-two-hybrid assays by conditional negative selection

**DOI:** 10.1101/2020.04.16.045757

**Authors:** Devendra Shivhare, Irene Julca, Pawel Gluza, Marek Mutwil

## Abstract

Yeast-two-hybrid (Y2H) is widely used as a strategy to detect protein-protein interactions (PPIs). Recent advancements have made it possible to generate and analyse genome-wide PPI networks en masse by coupling Y2H with next-generation sequencing technology. However, one of the major challenges of yeast two-hybrid assay is the large amount of false-positive hits caused by auto-activators (AAs), which are proteins that activate the reporter genes without the presence of an interacting protein partner. Here, we have developed a negative selection to minimize these auto-activators by integrating the *pGAL2-URA3* fragment into the yeast genome. Upon activation of the pGAL2 promoter by an AA, yeast cells expressing *URA3* cannot grow in media supplemented with 5-Fluoroorotic acid (5-FOA). Hence, we selectively inhibit the growth of yeast cells expressing auto-activators and thus minimizing the amount of false-positive hits. Here, we have demonstrated that auto-activators can be successfully removed from a *Marchantia polymorpha* cDNA library using *pGAL2-URA3* and 5-FOA treatment, in liquid and solid-grown cultures. Furthermore, since *URA3* can also serve as a marker for uracil autotrophy, we propose that our approach is a valuable addition to any large-scale Y2H screen.

## Introduction

The yeast two-hybrid (Y2H) assay is one of the most widely utilized approaches to identify protein-protein interactions (PPIs)(Yu *et al.*, 2011; Rolland *et al.*, 2014; Stark, 2006; Dreze *et al.*, 2011; Szklarczyk *et al.*, 2015; Lee *et al.*, 2015). Y2H data has been invaluable in revealing complexes regulating disease (Wang *et al.*, 2012), predicting gene function in plants (Dreze *et al.*, 2011), and revealing protein complexes underlying different plant-pathogen infections (Jiang *et al.*, 2016). However, genome-wide Y2H data generation is still challenging due to the cost and labor, needing to mate the specific constructs and to perform the necessary screening (Venkatesan *et al.*, 2009).

Recent approaches that combine next-generation sequencing (NGS) with Y2H are bringing us closer to obtaining genome-wide PPI networks (Weimann *et al.*, 2013; Yachie *et al.*, 2016). To avoid the manual, labor-intensive screening of bait proteins for tracking interactions, pools of baits are screened against pools of preys with multiplexed screening strategies (Yachie *et al.*, 2016; Hastie and Pruitt, 2007; Trigg *et al.*, 2017). Barcode Fusion Genetics (BFG-Y2H)(Yachie *et al.*, 2016) utilizes intracellular DNA recombination of barcoded open reading frame (ORF) to identify interacting proteins, thus allowing pooling and simultaneous sequencing of Y2H-positive colonies. Another approach, Cre-reporter-mediated Y2H coupled with next-generation sequencing (CrY2H-seq), uses Cre recombinase as a Y2H protein-protein interaction reporter. Cre covalently and unidirectionally links the interacting bait and prey plasmids via specialized loxP sites that flank the protein-coding sequences (Trigg *et al.*, 2017). The linked protein-coding sequences can be amplified by PCR and serve as interaction-identifying DNA molecules that can be sequenced by NGS approaches, thus allowing massively multiplexed screening for protein–protein interactions.

A common problem of the Y2H system is the autoactivation of Y2H-inducible reporter genes (Dreze *et al.*, 2010), which can result in a large number of false positives. This occurs when either DNA binding domain (BD) or activation domain (AD) activates transcription of Y2H reporter genes irrespective of the presence of any PPI. Three types of autoactivators (AAs) can be observed: (i) transcription factors that natively contain a functional AD that is active when fused to BD, (ii) proteins that are not transcription factors but contain a cryptic AD, and (iii) proteins that are not transcription factors, that contain one or more cryptic ADs that are only functional as truncated fragments.

Regardless of their type, AAs remain a significant hindrance in obtaining a successful PPI. In smaller screens, the bait proteins are typically screened for self-activation before commencing the assay, whereas, in experiments where thousands of bait proteins are tested for interactions with thousands of prey proteins, this might not be feasible. Furthermore, while it is possible to bioinformatically identify AAs (as they will appear to be interacting with an empty vector), the number of AAs tend to vastly exceed the numbers of true protein-protein interactions. For example, in the CrY2H study, 95% (164,293 out of 173,000) of all interactions were false positives caused by AAs (Trigg *et al.*, 2017). This indicates that only 5% of the generated data can be used in such assays.

To remove AAs, we have designed and introduced a *pGAL2-URA3* fragment to the yeast genome. *URA3* encodes orotidine-5-phosphate decarboxylase (ODCase), that can transform 5-fluoroorotic acid (FOA, a uracil analogue) into a highly toxic compound, 5-fluorouracil (FU)(Boeke *et al.*, 1984). Since *URA3* is driven by a pGAL2 promoter that is activated by AAs, the addition of FOA should selectively harm yeast cells containing AAs. We validated our system by identifying and removing AAs from *Marchantia polymorpha*, a model bryophyte.

## Materials and methods

### Nuclear transformation of yeast with pGAL2-URA3 fragment

To enhance the efficiency of the recombination, the DNA fragment containing the pRS305K-pGAL2-URA3 fragment and KanMX (kanamycin) selection marker was cleaved with AatII and KasI, gel-purified and used for yeast transformation. Yeast cells were inoculated from a fresh pre-culture and grown to OD600 of 0.5–0.7 at 28°C in YPD medium. The cells were harvested by centrifugation (3000 rpm for 3 minutes (min), 4°C), washed twice with 50 ml of ice-cold, sterile water, once with 50 ml of ice-cold, sterile 1M Sorbitol / 1mM CaCl2. The cells were harvested by centrifugation (3000 rpm for 3 minutes, 4°C) and resuspended gently in a 20 mL of ice-cold conditioning buffer (0.1M Sorbitol/10 mM DTT). The cells were kept for 30 min at 28°C with shaking (140 rpm) and then harvested by centrifugation (3000 rpm for 3 minutes, 4°C), washed with 50 ml of ice-cold, sterile 1M Sorbitol / 1mM CaCl2. The cells were resuspended in 2 mL of ice-cold, sterile 1M Sorbitol / 1mM CaCl2, and used immediately for electroporation. A total of 300 ul of prepared yeast cells were incubated on ice with 500 ng of gel-purified pGAL2-URA3 DNA cassette for 5 min. The cells were transferred to a 0.2 cm gap electroporation cuvette and electroporation pulse was applied at 2.5 kV. The efficient transformation resulted in electroporation time of 4-5 ms. The electroporated cells were immediately diluted with 2 ml of YPD medium, incubated on a shaker (650 rpm) for 4 hours (h) at 28°C and spread on YPD plates containing Geneticin (200 μg/mL). Plates were grown for 48 h at 28°C, and resistant colonies were transferred to a fresh selection plate and grown for 48 h at 28°C.

### Yeast strains and plasmids

CrY2H-seq yeast strains and AD and BD plasmids were obtained from Prof. Joseph Ecker’s lab, Salk Institute of Biological Sciences, La Jolla, USA. The strains and plasmids are also available from Arabidopsis Biological Resource Center (https://abrc.osu.edu/)(Trigg *et al.*, 2017). The two yeast strains were transformed by the pRS30SK-pGAL2-URA3 fragment and the resulting genotypes of the strains are:

- Y8800-URA3: *P*_*GAL2*_-*URA3 MATa leu2-3,112 trp1-901 his3-200 ura3-52 gal4Δ gal80Δ P*_*GAL2*_-*ADE2 LYS2::P*_*GAL1*_-*HIS3 MET2::P*_*GAL7*_-*lacZ cyh2*^*R*^
- CRY8930-URA3: *P*_*GAL2*_-*URA3 MATα leu2-3,112 trp1-901 his3-200 ura3-52 gal4Δ gal80Δ P*_*GAL2*_-*ADE2 LYS2::P*_*GAL1*_-*HIS3 MET2::P*_*GAL7*_-*CRE-HPHMX6 cyh2*^*R*^

### Yeast growth media

Yeast strains were grown in YEPD media, pH 6.0, containing 1% yeast extract, 2% peptone, 200 mg/L adenine hemisulfate and 2% agar (only for solid) supplemented with 50 mL of 40% glucose and antibiotic, G418 (200 μg/mL) and G418 + hygromycin (125 μg/mL) for yeast strains Y8800 and CRY8930, respectively

### Yeast selection media

The selection for yeast strains harboring pDEST_AD_lox_TRP1 or pDEST_DB_lox_LEU2 plasmids was done using synthetic complete (SC) drop-out media containing 6.7 g/L yeast nitrogen base, 1.47 g/L (minus adenine (A), histidine (H), leucine (L), tryptophan (T)) or 1.37 g/L (minus AHLT and minus uracil) dropout mix, 200 mg/L adenine hemisulfate and 2% agar (only for solid) supplemented with 50 mL of 40% glucose, 8 mL/L of desired amino acid solution (100 mM leucine/histidine and 40 mM tryptophan stock solution). For SC-FOA media, 0.2 % 5-FOA (5-Fluoroorotic acid hydrate, 100 mg/mL stock in DMSO) was added after autoclaving the media. The selection media was additionally supplemented with G418 (200 μg/mL) to avoid microbial contamination.

### Growth conditions for *Marchantia polymorpha*

Tissue culture grown *Marchantia polymorpha* was obtained from Prof. Chen Zhong from NIE, Singapore. The plant was propagated in the lab using half Gamborg B5 media without sucrose (pH 5.5, 1.4% agar) on deep tissue culture plates under 24 hours light and 22-23 °C.

### RNA extraction and construction of entry cDNA library for *M. polymorpha*

The total RNA was extracted from 2-3 weeks old thallus using ReliaPrep Promega kit as per manufacturer’s protocol. The integrity of the RNA samples was confirmed using bioanalyzer (Agilent 2100). The samples that passed quality control (RIN>8.5) were pooled together, concentrated using Zymo RNA concentrator kit, and rechecked for the RIN value.

cDNA was synthesised, and the entry library was prepared using CloneMiner™ II cDNA library construction kit (Invitrogen) using Gateway technology, as per manufacturer’s instruction with slight modifications. Briefly, the total RNA was used as an mRNA template, and reaction was primed by hybridizing Biotin-attB2-Oligo(dT) to the poly(A) tail, followed by synthesis of first strand using SuperScript III Reverse transcriptase provided with the kit. The second strand was synthesized using the first strand as a template. The synthesized cDNA was extracted using phenol:chloroform:isoamyl alcohol (25:24:1) and precipitated using ethanol before finally dissolving in DEPC-treated water. The 5’ end of the cDNA is ligated with attB1 Adapter followed by size fractionation using columns provided with the kit to eliminate residual adapters and other low molecular weight DNA. Subsequently, the BP recombination reaction was performed using Gateway® BP clonase® II enzyme mix to facilitate recombination of attB-flanked cDNA into pDONR 222™ entry vector. After that, the reaction was inactivated with proteinase K, and the cDNA was precipitated using ethanol. The cDNA pellet was resuspended in TE buffer.

The resuspended cDNA was divided into six aliquots and each aliquot was transformed into ElectroMax™ DH10B™ T1 phage resistant competent cells in a separate vial using a Gene pulsor Xcell™ electroporation system (Bio-Rad) as mentioned in the kit’s manual. After transformation, all the vials were pooled together, further aliquoted, and stored at −80°C.

### Analysis of the cDNA entry library

A small sample from the above library was utilised to perform the plating assay for titre determination as per kit’s manual. The sample was serial diluted and plated in duplicates on LB plates with kanamycin selection marker. Next day, colonies were counted and the titre for each plate was determined using the equation below:

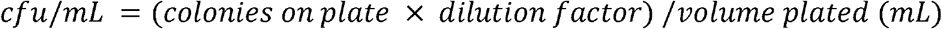

The titre from each plate was used to calculate the average titre for the whole cDNA library. The total colony forming units (CFU) were determined using the following equation:

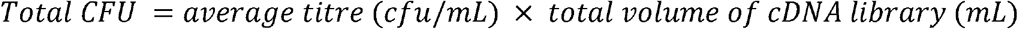

The cDNA library was further qualified to determine the average insert size by setting up colony PCR reactions using M13 forward and reverse primers. We picked a total of 19 colonies and one vector control. The PCR reaction was set using Exten 2x PCR master mix (first base) as per manufacturer’s protocol.

### Generation of yeast destination expression library

The cDNA library (entry) was shuttled into the Gateway® destination vector (pADlox or pDBlox) to generate expression clones/library using the LR recombination reaction with Gateway® LR Clonase® II enzyme mix as per manufacturer’s protocol. The titre determination of each library was done using plating assay, as previously mentioned.

### Preparation of electro-competent cells and transformation of cDNA expression libraries into yeast strains using electroporation

One vial of the glycerol stock from each destination expression library was thawed, incoluted in LB + Carbenicillin (100 μg/mL) and grown overnight at 37 °C at 220 rpm for plasmid isolation. Next day, plasmid DNA was isolated using Wizard® *Plus* SV Minipreps purification kit from Promega and used for yeast transformation by electroporation as described in (Benatuil *et al.*, 2010) with modifications.

Yeast strains, Y8800, and CRY8930 were streaked out on YEPD plates supplemented with G418 (200 μg/mL) and G418 + hygromycin (125 μg/mL), respectively, and incubated at 37°C for 48 hours. For each strain, 10-12 colonies were picked from the plates and grown in 20 mL of YEPD overnight at 30 °C and 220 rpm to attain a stationary phase. The next day, the OD600 was measured, and the cells were diluted up to a total volume of 100 mL of YEPD, aiming for an initial OD600 between 0.1-0.2. The cells were further grown until the OD600 reached approximately 1.0-1.5 and underwent at least two doublings, which usually takes 5-6 hours. Yeast cells were collected by centrifugation at 3000 rpm for 3 minutes, media was removed, and pellet was washed twice with 50 mL ice-cold water and once with 50 mL of ice-cold electroporation buffer (1 M Sorbitol / 1 mM CaCl_2_). Yeast cells were conditioned by re-suspending the cell pellet in 20 mL 0.1 M Lithium acetate/10 mM DTT and shaking at 220 rpm in a culture flask for 30 minutes at 30 °C.

The conditioned cells were collected by centrifugation, washed once by 50 mL ice-cold electroporation buffer, and re-suspended in 100 to 200 μL electroporation buffer to reach a final volume of 1 mL. The electrocompetent cells were now ready to use and kept on ice until electroporation.

The electroporation reactions of 400 μl each were set by gently mixing electrocompetent cells with a library plasmid DNA. The reaction mixture was transferred to a pre-chilled electroporation cuvette (0.2 cm width, BioRad) and kept on ice for at least 5 mins before electroporation. The cells were electroporated at 2.5 kV and 25 μF and immediately supplemented with 1 ml of 1:1 mix of 1M sorbitol:YEPD media. The entire content from each cuvette was further transferred to an additional 8 mL of 1:1 mix of 1 M sorbitol:YEPD media as soon as possible and incubated for 1 h at 30 °C and 220 rpm. Cells were collected, washed with sterile water once, and finally resuspended in 10 mL SC-Leu and SC-Trp media for bait and prey strains, respectively. An aliquot from each transformed cell mixture was serial diluted (1/100, 1/1000) and plated on SC-Leu and SC-Trp plates for bait and prey strain respectively and incubated at 30 °C. The remaining cells were grown for 2 days at 30 °C and 220 rpm, glycerol stocks were prepared, aliquoted, and stored frozen at −80 °C. The size of each library was determined from the colony counts after 3 days. This was the final bait and prey transformed library.

### PCR amplification of the transformed library and Next-generation sequencing (NGS)

One vial each of transformed bait and prey library was thawed and inoculated into 10-12 ml of SC-Leu and SC-Trp, respectively, at 30 °C and 220 rpm overnight, with the corresponding antibiotics.

The next day, yeast cells were collected, washed, and resuspended in 0.5-1 mL of yeast lysis buffer (50 mM Tris-HCl pH 7.5, 0.92 M sorbitol, 10 mM EDTA and 1% β-mercaptoethanol). Additionally, 200 U of lyticase (Sigma) was added to each tube to digest the cell wall by incubating at 37 °C for 1-2 hours with mild shaking (200 rpm). After incubation, yeast cells were collected by centrifugation (3000 rpm, 5 mins), and the supernatant was discarded. The plasmid DNA isolation was performed with Wizard® *Plus* SV Minipreps DNA Purification System as per the manufacturer’s protocol. The isolated plasmids were used for PCR amplification of the individual library with specific forward (AD:5’ CACTGTCACCTGGTTGGACGGACCAAACTGCGTATAACGC or BD: 5’ GATGCCGTCACAGATAGATTGGCTTCAGTGG) and reverse (Y2H term reverse: 5’ GGAGACTTGACCAAACCTCTGGCG) primers using Phusion High-Fidelity PCR Kit from Thermo Scientific, with a standard PCR program.

The multiple PCR reactions were set up for each library, pooled together, and the PCR product was loaded on a 1% agarose gel in TAE buffer and ran at 100 V for 40-50 mins. The PCR product >500 bp was excised and purified from the gel using Wizard® SV Gel and PCR Clean-Up System as per kit’s instruction. The purified PCR product was sent to NovogeneAIT (https://en.novogene.com/) for sequencing. The platform used was NovaSeq 6000 - Illumina, and 6 Gb / sample of paired-end reads of 150 nt and insert size <= 500bp was generated. Finally, to determine the gene coverage of the individual library, the reads were mapped to Marchantia genome (Bowman *et al.*, 2017) using Diamond v0.9.25 (Buchfink *et al.*, 2014). Reads that mapped to the coding sequences of Marchantia with significant (e-value<0.001) value were deemed to indicate that the matching coding sequence is present in the library. The raw reads for the activation domain (AD) and binding domain (BD) library are available at the ENA archive at ERRXX (Marek: the fastq files are on a hard drive in my office, and I will upload the fastq files once we can get back to work. My apologies).

### Generating auto-activator control for BD library

One vial of the bait library (BD) was thawed and plated on multiple SC-Leu and SC-HULA plates to determine the average percentages of auto-activators in our library. After 3 days, the colonies were counted, and SC-HULA plates were scraped, resuspended in SC-Leu media, and glycerol stocks were made to create archival BD auto-activator library. A vial from −HULA library was thawed, serially diluted, and plated on −HULA plates. After 3 days, five independent colonies were picked and incubated in liquid −LH+G418 media overnight at 30 °C. Glycerol stocks were prepared from the overnight culture as representatives of auto-activators of BD library. Also, five overnight cultures were serially diluted (0, 1:10, and 1:100) and spotted on −L, −L+FOA, -HULA plates for further validation. The auto-activators genes were characterized by Sanger sequencing, where isolated plasmids/PCR fragments were sent for forward sequencing by using forward BD primer. After sequencing, the query sequences were matched with pDBlox plasmid sequences using MultAlin multiple sequence alignment web tool (Corpet, 1988).

### Removal of auto-activators by 5-FOA in liquid cultures

The five auto-activators (AAs) screened above, and three empty vector (EV) colonies (negative control) were grown individually in different media (SC-Leu, SC-Leu+FOA, and SC-LH) for three days. OD600 was measured every 24 hours in a 96-well plate to test the influence of the media on growth using a microplate reader.

### Removal of auto-activators by 5-FOA in solid media

Five AAs and one empty vector strain were grown individually overnight in SC-Leu media. The initial ODs of the overnight grown cultures were set to 0.001 each, the cultures were mixed to achieve the final OD of 0.006 (0.001 each for 5 AAs and 1 empty vector (EV)) and grown overnight in SC-Leu media. The next day, the mixed culture was inoculated in SC–Leu (selection for BD plasmid only), SC-Leu+FOA (to remove AAs), and SC-Leu-His (to remove non-AAs) for 24 hours. The next day, the cultures from the SC–Leu, SC-Leu+FOA, and SC-Leu-His plates were plated on individual SC–Leu plates. After 2 days, 48 colonies from each plate were grown in liquid SC–Leu media overnight, and finally spotted (1:10 dilution) on −Leu+FOA and −Leu-His plates and observed after three days (Figure 3A).

## Results

### Designing conditional negative selection in yeast-two-hybrid with URA3

In our pursuit to develop a genetic tool that can remove auto-activators from the yeast two-hybrid assay, we had to establish two components of the tool. First, we needed a negative selection marker that can inhibit the growth or kill the yeast cells. Second, we needed a means to conditionally activate the negative selection marker only when an auto-activator was present in the yeast cell.

To identify a suitable negative selection marker, we evaluated ten markers available for yeast (Siewers, 2014). The most well-established negative selection marker, *URA3*, has been used for decades in positive and negative selection in *Saccharomyces cerevisiae*. *URA3* encodes orotidine-5-phosphate decarboxylase (ODCase), which can catalyze the transformation of 5-fluoroorotic acid (FOA, a uracil analogue) into a highly toxic compound, 5-fluorouracil (FU)(Boeke *et al.*, 1984). Thus, the activity of *URA3* confers sensitivity to FOA, and cells not expressing *URA3* are FOA resistant. Alternatively, *URA3* can be used as a positive selection marker, as it can complement uracil auxotrophy of a *URA3*-negative strain. To implement the conditional activation of *URA3*, we assessed systems that would induce the expression of *URA3* only when an auto-activator is present. Since auto-activators induce expression of positive selection makers (His, Ade) that are driven by pGAL-family promoters, we chose to use pGAL2 promoter to drive *URA3*.

We synthesized pGAL2-URA3 fragment and cloned it into pRS305K plasmid, which can be used to integrate the fragment into Leu2 chromosomal locus and select for the transformation by using kanamycin (Figure 1A). The sequence of the vector is available as Supplemental Data 1. We have integrated the fragment into the genomes of yeast strains used in CrY2H-seq (Trigg *et al.*, 2017), which use a Cre recombinase to fuse *in vivo* the coding sequences of two interacting proteins, allowing to identify these interactions with next-generation DNA sequencing.

**Figure 1.**
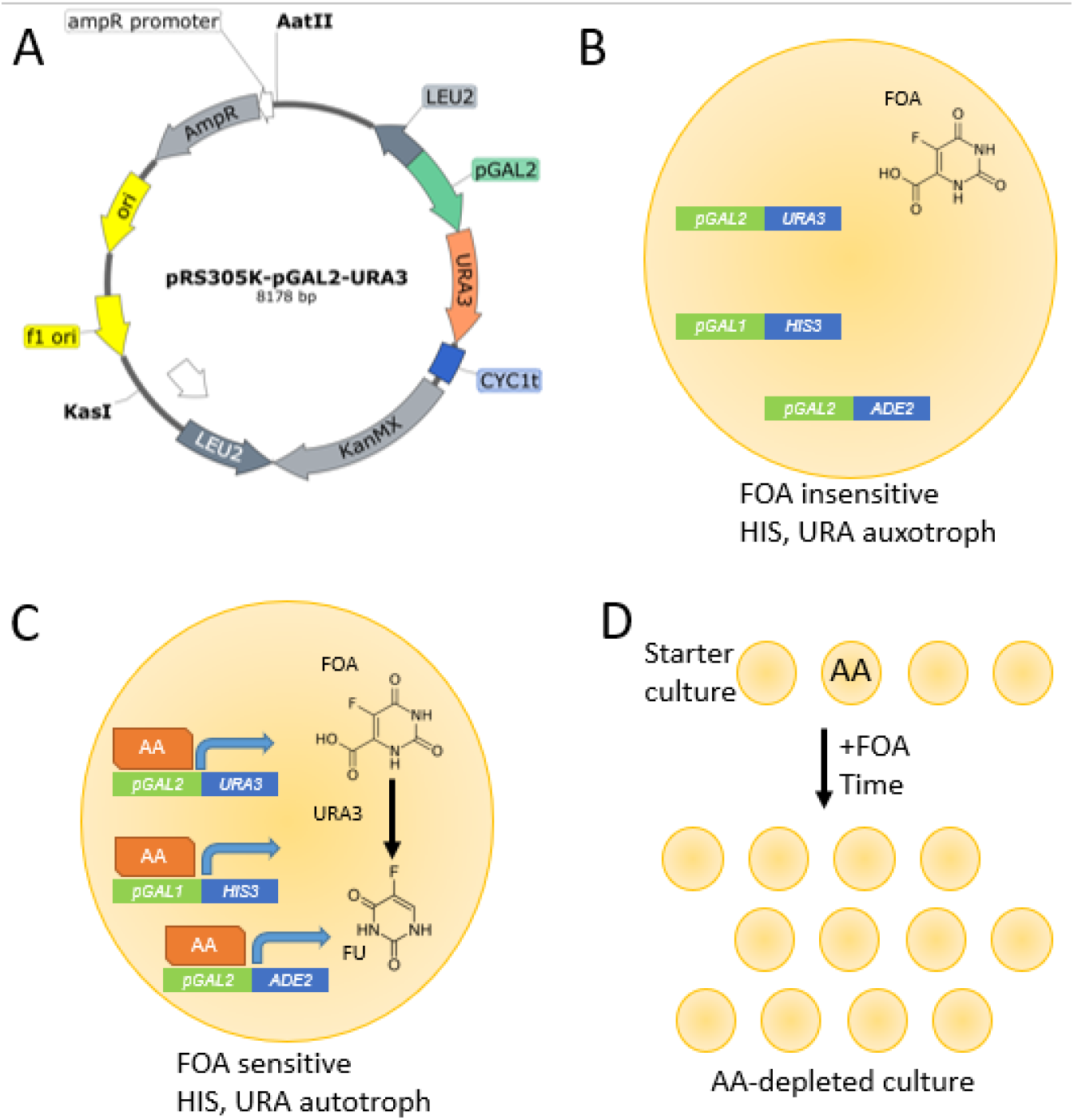
Synthesized pGAL2-URA3 plasmid and outline of the method. **(A)** pRS305k-pGAL2-URA3 plasmid harboring *URA3* driven by pGAL2 promoter. (B) Scenario where the yeast cell does not contain an auto-activator (AA). *URA3* and *HIS3* are not expressed, and FOA is not converted into FU. C) Scenario where an AA activates the expression of *URA3* and *HIS3*, resulting in the conversion of FOA into toxic FU by *URA3*. D) Description of the AA-removal method, where a starter culture containing AAs is treated by FOA. Since FOA can inhibit or kill yeast, the growth of AAs should be severely delayed or outright stopped by FOA.

A yeast strain containing this new fragment in the presence of FOA, but with the absence of an auto-activator, would not express *HIS3* and *URA3* and would be FOA insensitive and histidine auxotrophic (Figure 1B). Conversely, in the presence of an auto-activator, both *HIS3* and *URA3* would be expressed, and the yeast would be histidine and uracil autotroph and sensitive to FOA (Figure 1C).

We hypothesize that by adding FOA to the media before the mating, growth of haploid yeast cells that express *URA3* should be inhibited (Figure 1D), due to the accumulation of the toxic FU in the yeast cells. This should reduce or eliminate the number of auto-activators in our library population, and subsequently reduce the number of false-positive hits when used for mating in a yeast two-hybrid assay.

### Construction of AD and BD libraries for *Marchantia polymorpha*

To test our AA-removal method, we set to identify genes that are AAs in the activation domain (AD) and the DNA binding domain (BD) libraries. We chose *Marchantia polymorpha* as a source for AAs, as it is an emerging model for early diverging plants, has a low number of genes, and minimal genetic redundancy (Bowman *et al.*, 2017).

The cDNA library was successfully generated from *Marchantia* thallus mRNA, moved into *E. coli*, where a plate assay revealed that 6.276*10^5^ entry clones were produced. Colony PCR analysis of 19 individual colonies confirmed inserts of variable size ranging from >1 kb to <3kb (Figure S1). Following this, the entry library was transferred into destination vectors using LR recombination reaction to generate expression clones. The final titre of the cDNA expression library for *M. polymorpha* was determined to be 5.48×10^5^ and 6.12×10^5^ for AD and BD clones, respectively.

The expression libraries were successfully electro-transformed into their respective host yeast strains. The plating assay performed with transformed libraries revealed a total of 3.4×10^5^ and 2.86×10^5^ transformants from AD and BD libraries, respectively.

To estimate the percentage of *Marchantia* coding sequences that are captured by our AD and BD libraries, we amplified the cloned CDSs by PCR, which revealed an expected smear on the gel (Figure S2). The purified DNA was subjected to Illumina sequencing, which revealed that 7,325 and 7,035 genes were present in AD and BD libraries, respectively, while 5,926 genes were present in both. This implies that our libraries contain ~1/3 of the total CDSs, indicating reasonably good coverage.

### Quantification of AAs in AD and BD libraries

To quantify the amount of AAs in AD and BD libraries, we used a plate assay to compare the number of yeast colonies that grow on media selecting for activation of *His* (H), *Ura* (U) and *Ade* (A) markers (Figure 1C), versus the total number of colonies. BD library was grown on −L (selecting for presence of pDBlox plasmid) and -HULA (additional selection for expression of *His*, *Ura* and *Ade* markers), while AD library was grown on -Trp (T, selecting for presence of pADlox plasmid) and -HUTA (additional selection for *His*, *Ura* and *Ade* markers). After 3 days, the colonies were counted, and the average percentages of auto-activators were calculated by dividing the number of colonies on -HULA plates against the number of colonies on −L plates.

We found that the BD library contains 0.13% (448 colonies on -HULA plates, 345000 colonies on −L plates) auto-activators (Figure 2A). Conversely, we rarely found any AD library colonies on -HUTA plates, and only 0.0029% (13 colonies on -HUTA plates, 484000 colonies on -T plates) of auto-activators were found (Figure 2A).

**Figure 2.**
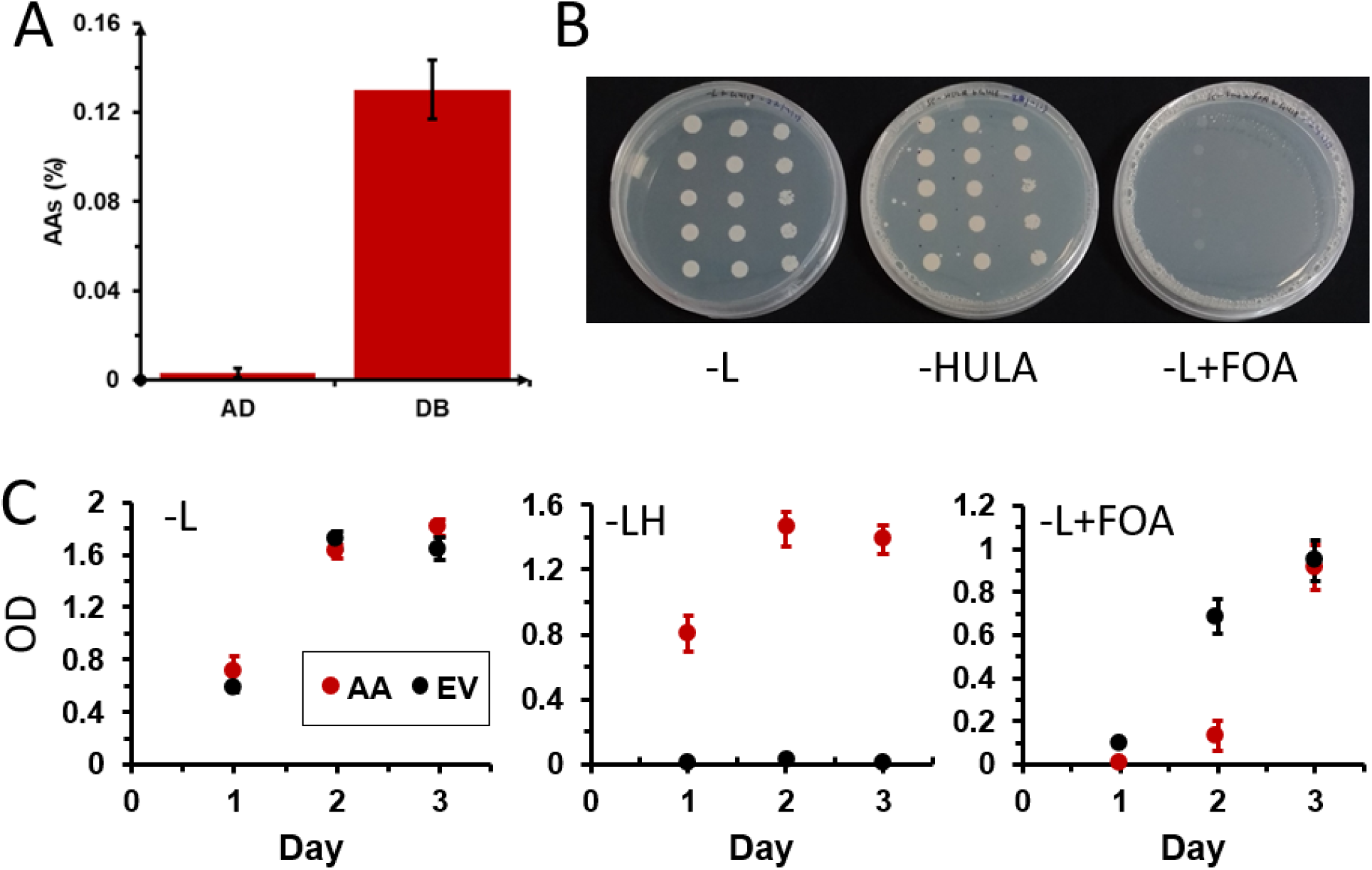
Quantification of auto activators in AD and BD and removal by 5-FOA treatment in liquid cultures. (A) Percentage of the auto-activators present in AD and BD libraries. The error bars indicate standard deviation. (B) Growth of the five AAs from the BD library on −L (left), -HULA, and −L+FOA (right). AA1-5 (rows) were spotted in three serial dilutions (1:1, 1:10, 1:100, columns) and grown for 3 days. C) Growth of the five AAs from the BD library (red points) and three empty vector (EV, black points) controls in liquid −L (left), −LH, and −L+FOA (right).

**Figure 3.**
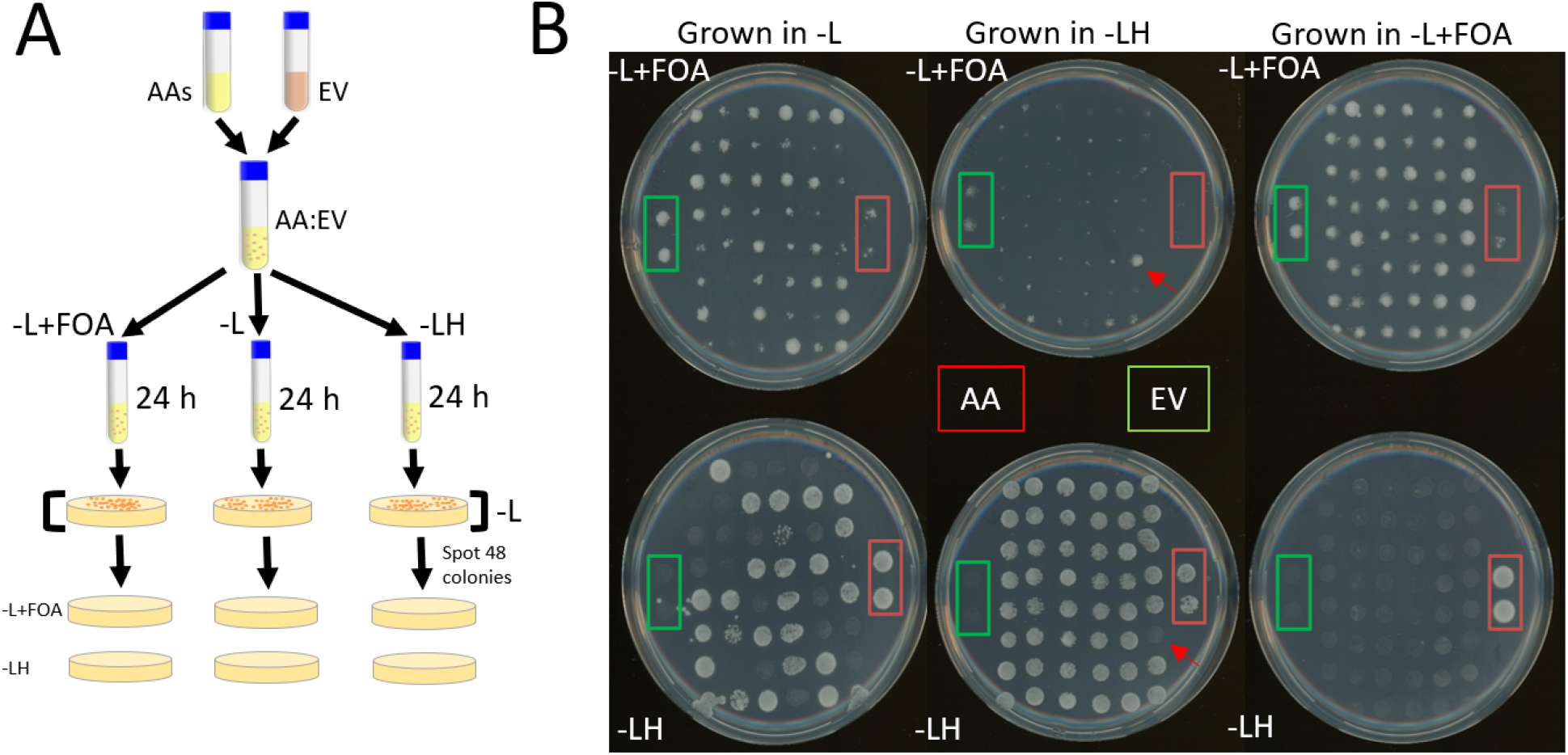
Removing auto-activators with FOA. (A) The experimental design. The AAs and EV (non-AAs) were mixed. The next day, the mixed culture was inoculated individually in −Leu (no selection), −L+FOA (putatively removing AAs), and −LH (putatively removing non-AAs). The cultures were plated on −L plates and 48 colonies were picked and grown in −L media. The 48 colonies were parallel-spotted on −L+FOA and −LH plates. (B) Growth of −L-grown AA:EV culture (left column), on −L+FOA plate (top), and −LH (bottom). The middle column contains colonies passed through −LH, while the right column contains colonies passed through −L+FOA. The two colonies marked with red and green boxes indicate AAs and EV, respectively.

To reveal the identities of the AAs from the BD library, we chose five colonies (AA1-AA5) growing on -HULA plates and subjected them to Sanger sequencing. Subsequent BLAST analysis against *Marchantia polymorpha* genome revealed that AA1 best matched with a hypothetical protein MARPO_0107s0034 (putative histone deacetylase), AA2 matched MARPO_0055s0078 (putative ubiquitin-protein ligase), AA3 and AA4 both matched MARPO_0101s0042 (putative NEDD8, ubiquitin-dependent protein catabolic process) and AA5 matched AXG93_4751s1100 (putative 40S ribosomal protein)(Table S1).

Since AAs were more frequent in the BD library, we decided to use the five colonies growing on -HULA as a validation of our AA-removal method.

### FOA can suppress the growth of AAs on solid and liquid media

To test whether FOA can inhibit the growth of AAs, we plated the five AAs on solid media containing −L (selection for pDBlox plasmid), -HULA (selection for AAs), −L+FOA (0.2 % 5-FOA). As expected for the AAs, we observed the growth of AA1-5 on −L and -HULA (Figure 2B). Conversely, we did not see any growth on FOA plates, suggesting that FOA suppressed the growth of the AAs.

To further confirm this result and see whether this system applies to liquid-grown cultures, we have compared growth rates of AAs, and empty vector (EV) controls in −L, −LH and −L+FOA. In −L media, since there was no positive or negative selection for the pGAL2-activated genes, the auto-activators (AA, red) and empty vector strains (EV, black) showed similar growth pattern for all three days (Figure 2C, left). On the other hand, in the presence of −LH media, as expected for a scenario where pGAL-driven genes are selected for, AAs grow normally, reaching saturation on the second day, while EV did not show any significant growth even after 3 days (Figure 2C, middle).

However, when SC-Leu media was supplemented with 0.2% FOA, AAs did not show observable growth after 1 day, while the EV showed a slower growth (Figure 2C, right). On day 2, AAs showed minor growth, and their OD was 5.3 times lower compared to EV, indicating that the growth of AAs is penalized in FOA. On day 3, auto-activators and EV showed similar growth. This indicates that FOA can inhibit the growth of AAs, but the negative selection loses its potency after day 2 in liquid media.

### Removal of auto-activators from a mixed culture

After demonstrating the inhibiting effect of FOA on AAs grown in solid and liquid culture, we investigated whether we can remove AAs from a mixture of yeast cells containing AAs and non-AAs (EV, empty vector controls). To this end, we have mixed AA:EV liquid cultures and subjected the resulting mix to no selection (−L), enrichment for AAs (−LH), and removal of AAs (−L+FOA). After 24 hours of incubation, 48 colonies from the three cultures were parallel-spotted on plates selecting non-AAs (−L+FOA) or selecting AAs (−LH, Figure 3A). As a control, we have spotted two colonies of AAs (red square) and non-AAs (green square, Figure 3B).

The AA:EV culture that was passed through −L, which imparts no positive or negative selection, we observed 26 colonies growing on the −LH plate, indicating that roughly half of the 48 colonies are AAs (Figure 3B, left-bottom plate). As expected, AA controls (red square) grew well on −LH, but poorly on −L+FOA, while non-AA control showed an opposite behavior (green square).

Growing the AA:EV culture in −LH should have resulted in a culture composed predominantly by AAs. Indeed, 47 out of 48 colonies were growing well on −LH (Figure 3B, middle-bottom plate), but poorly on −L+FOA (Figure 3B, middle-top plate). Interestingly, the one colony growing poorly on −LH (red arrow), grew well on −L+FOA, indicating that this colony is a non-AA that escaped the −LH selection. Finally, the tested whether the AA:EV culture grown in −L+FOA was depleted of all AAs. Indeed, we observed no colonies on −LH (Figure 3B, bottom-right), indicating that our method has successfully removed AAs from the mix.

Taken together, based on results from this and previous sections, we can conclude that the negative selection using pGAL2-URA3 and FOA screening works well in liquid and solid media, preventing the growth of AAs.

## Discussion

Since AAs are a large problem in Y2H screens and are difficult to remove by screening when performing a large-scale assay where millions of interactions are tested, we have developed a method that removes AAs by using conditional negative selection with *URA3* gene, driven by pGAL2 promoter. The pGAL2 promoter activates *URA3* only in the presence of an AA or a genuine PPI, allowing *URA3* to convert 5-FOA into a toxic FU. When applied to haploid AD and BD cultures, the method has thus potential to selectively minimize or outright eliminate yeast cells harboring AAs.

The previous study has shown that AAs comprise 16% of baits (Trigg *et al.*, 2017), which in turn comprised 95% of detected interactions in their large-scale CrY2H assay. In our study, we observed that BD and AD libraries contained 0.13% and 0.0029% AAs, respectively (Figure 2A). We speculate that the higher number of AAs observed in the CrY2H study is because the authors focused on transcription factors (Trigg *et al.*, 2017), which are likely to contain bona fide activation domains. Conversely, since we have studied all coding sequences in *Marchantia polymorpha*, the proportion of transcription factors containing activation domains should be lower, explaining the lower number of observed AAs. Despite the lower numbers in Marchantia, removal of AAs is still a worthwhile pursuit as it will result in a higher number of detectable true positive interactions.

We observed that the FOA treatment effectively suppresses the growth of AAs on solid media (Figure 2B and 3B), and liquid media (Figure 2C). However, the efficacy of the assay in liquid media drops after three days of incubation, as the growth of AAs equals that of the non-AAs (Figure 2C, right panel). This indicates that our approach is only effective in the first two days. While the molecular basis for the loss of efficacy is unclear, we speculate that the unwanted growth could be caused by spontaneous mutations of the *pGAL2-URA3*, which either inhibit the activation of *URA3* or inactivate the enzyme. Further work should attempt to introduce another copy of pGAL2-URA3 into the genome to provide a genetic backup for the assay.

We were able to remove all AAs from the panel of 48 colonies, indicating that despite the loss of efficacy after two days of cultivation in FOA, the assay is still able to deplete the AAs from a yeast culture (Figure 3B). Furthermore, since the *URA3* gene is also activated by true interactions (Figure 2B), including pGAL2-URA3 into Y2H assays will have a valuable dual role of removing AAs and selecting for true PPIs.

## Supplementary Materials

### Supplementary Figures

**Figure S1.**
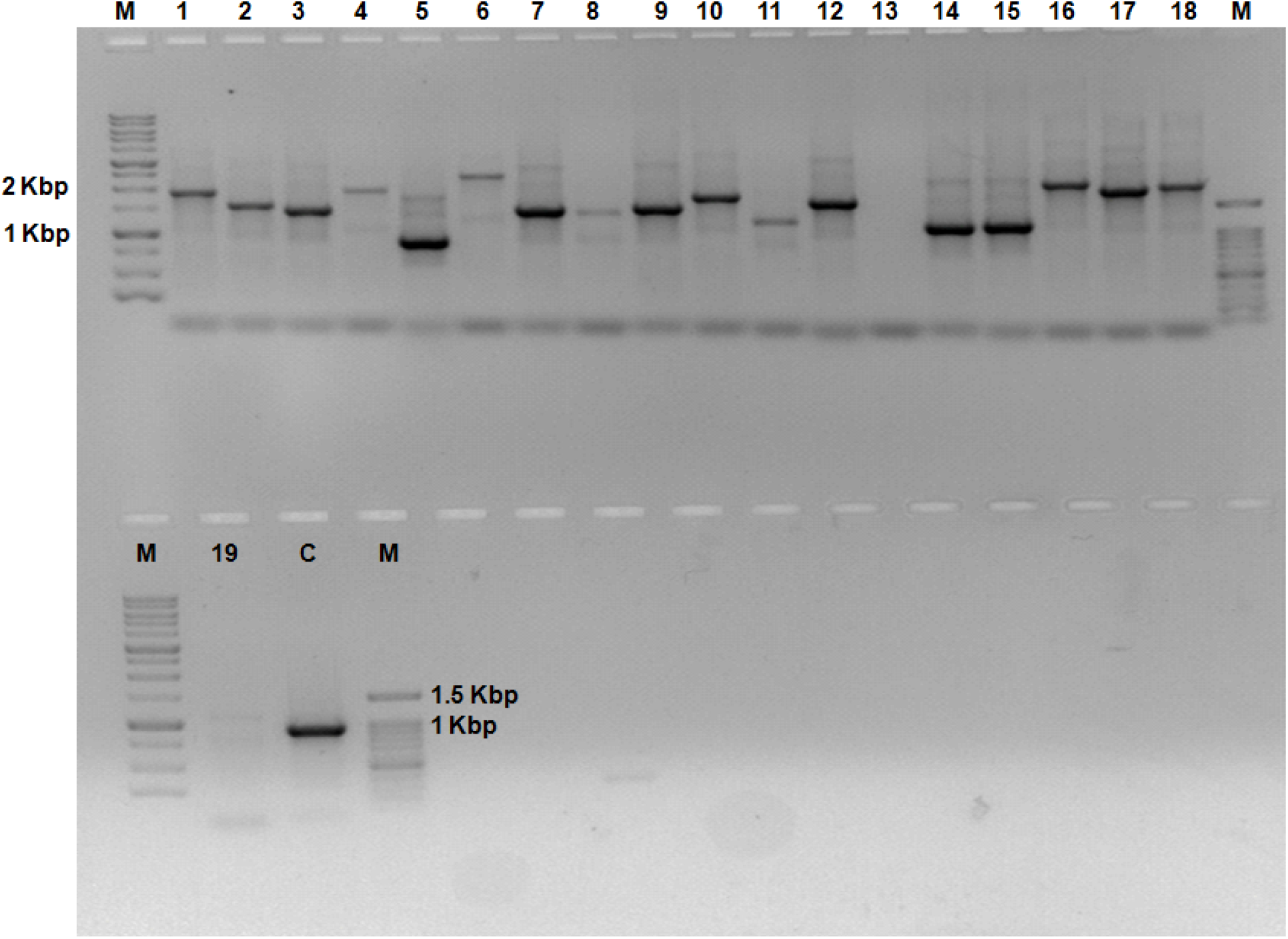
PCR analysis of 19 colonies cloned into the entry vector. The quality of the library was assessed by picking a total of 19 colonies for colony PCR run with the M13F+R primers. The PCR reaction was loaded onto 1% agarose. M: marker, 1-19: colonies, C: pDONR vector control.

**Figure S2.**
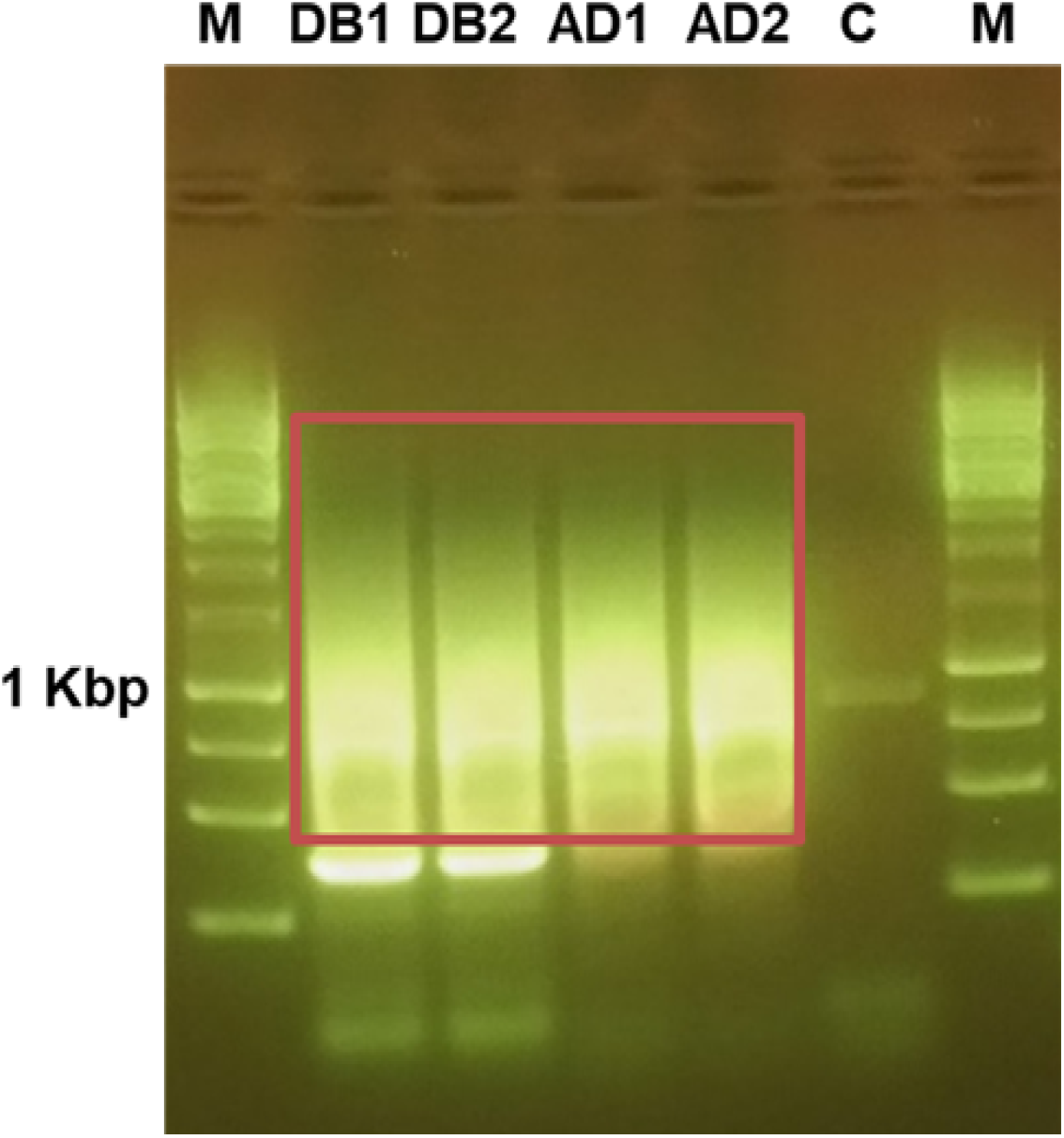
The plasmids were isolated from BD and AD libraries and PCR-amplified. The smear that was used to purify DNA from the gel and sent for NGS analysis is indicated by the red box.

### Supplemental Tables

**Table S1. Best BLAST hits of the five AAs to the Marchantia proteome.**

### Supplemental data

**Supplemental Data 1. pRS305K-pGAL2-URA3 plasmid in .dna format.**

## Author contributions

Conceptualization: M.M.; Experiments: D.S., M.M-L., P.G., NGS data analysis: M.M., I.J.

## Funding

This article was supported by Max Planck Gesellschaft and is funded by NTU Start-Up Grant and Singaporean Ministry of Education grant MOE2018-T2-2-053.

## Acknowledgments

We would like to thank Prof. Joseph Ecker for his kind gift of the vectors and yeast strains and Prof. Chen Zhong for giving us the Marchantia culture.

## Conflicts of Interest

## Notes

### Competing Interest Statement

The authors have declared no competing interest.

